# Intraoral thermal processing in the gustatory cortex of awake mice

**DOI:** 10.1101/2023.02.06.526681

**Authors:** Cecilia G. Bouaichi, Katherine E. Odegaard, Camden Neese, Roberto Vincis

**Author notes:** These two authors contributed equally to this work (Cecilia G. Bouaichi, Katherine E. Odegaard).

## Abstract

Oral temperature is a sensory cue relevant to food preference and nutrition. To understand how orally-sourced thermal inputs are represented in the gustatory cortex (GC) we recorded neural responses from the GC of male and female mice presented with deionized water at different innocuous temperatures (14 °C, 25 °C, 36 °C) and taste stimuli (room temperature). Our results demonstrate that GC neurons encode orally-sourced thermal information in the absence of classical taste qualities at the single neuron and population levels, as confirmed through additional experiments comparing GC neuron responses to water and artificial saliva. Analysis of thermal-evoked responses showed broadly tuned neurons that responded to temperature in a mostly monotonic manner. Spatial location may play a minor role regarding thermosensory activity; aside from the most ventral GC, neurons reliably responded to and encoded thermal information across the dorso-ventral and antero-postero cortical axes. Additional analysis revealed that more than half of GC neurons that encoded chemosensory taste stimuli also accurately discriminated thermal information, providing additional evidence of the GC’s involvement in processing thermosensory information important for ingestive behaviors. In terms of convergence, we found that GC neurons encoding information about both taste and temperature were broadly tuned and carried more information than taste-selective only neurons; both groups encoded similar information about the palatability of stimuli. Altogether, our data reveal new details of the cortical code for the mammalian intraoral thermosensory system in behaving mice and pave the way for future investigations on GC functions and operational principles with respect to thermogustation.

## Introduction

The consumption of food and beverages is highly dependent on the initial sensation and the response it evokes (Dotson et al., 2012; Schier and Spector, 2019). This sensation arises within the mouth and involves the integration of intraoral gustatory, olfactory, and somatosensory cues in a single percept called flavor (Kemp and Beauchamp, 1994; Small, 2012). In the past decades, many electrophysiological studies in behaving rodents have investigated the physiological correlates of one of these intraoral sensory components – taste – which originates when chemical compounds stimulate specialized chemoreceptors within the oral cavity (Vincis and Fontanini, 2019; Spector and Travers, 2005). Using gustatory stimuli at a fixed temperature, these studies made clear that taste information is processed through neural computations that occur in interconnected brain areas that include the gustatory cortex (GC), the primary cortical area responsible for processing taste information (Katz et al., 2001; Jezzini et al., 2013; Stapleton et al., 2006; Liu and Fontanini, 2015; Samuelsen et al., 2013; Roussin et al., 2012; Bouaichi and Vincis, 2020; Levitan et al., 2019). In addition, these studies have indicated that GC neurons respond to compounds representing different taste qualities and their hedonic value with time-varying patterns of activity (Katz et al., 2001; Jezzini et al., 2013; Bouaichi and Vincis, 2020; Arieli et al., 2020; Neese et al., 2022) and play a role in driving taste-related decisions (Vincis et al., 2020; Mukherjee et al., 2019; Vincis and Fontanini, 2016b).

However, a growing body of experimental work indicates that neurons in the GC are also capable of responding to non-gustatory components of intraoral stimuli (Bouaichi and Vincis, 2020; Stapleton et al., 2006; Rudenga et al., 2010; Samuelsen and Fontanini, 2017; De Araujo et al., 2003; Samuelsen and Vincis, 2021; Small et al., 2004; Maier, 2017), including temperature - a salient feature of the sensory properties of foods and beverages. Different studies in humans and primates (Verhagen et al., 2004; Cerf-Ducastel et al., 2001; Guest et al., 2007), as well as pioneering works in anesthetized rats (Yamamoto et al., 1981; Kosar et al., 1986), have indicated that changes in intraoral temperature seem to modulate activity in GC neurons. While these data implicate the GC as a potential key cortical region for the integration of taste and thermal orosensory inputs, they stop short of supplying a fine-grained analysis of the GC’s neural responses, and many questions remain. Here, using extracellular recording (tetrodes and silicon-based probes), we aim to provide a complete neurophysiological assessment of how thermal orosensory inputs shape GC activity in alert mice. Specifically, this study is designed to assess 1) whether and how neurons in the GC of actively licking mice are modulated by changes in the temperature of a chemically inert drinking solution, 2) the spatial organization of intraoral thermal-related information across the GC, and 3) the extent to which GC neurons respond to both intraoral thermal and taste information.

Our data provide novel insights into how thermal information originating from the mouth is integrated in the neurons of a central structure associated with taste. Our findings demonstrate that more than half of the neurons recorded throughout the GC encode deionized water information in a temperature-selective and mostly monotonic manner. Additionally, we show these responses to thermal stimuli occur in the absence of any overt taste quality; neural responses to deionized water and artificial saliva presented at the same temperature revealed no significant differences. Further analysis of our data suggests that thermal stimulation inside the mouth evokes responses that are organized in a coarse topographic manner along the dorso-ventral axis of the GC and as an antero-postero gradient. Our study finds that more than half of taste-responsive neurons are also capable of representing intraoral thermal information. While these neurons responding to both taste and temperature are shown to have broad tuning profiles, they do not appear to encode any more information regarding palatabilty than GC neurons responsive only to taste. Overall, our results are consistent with the GC being an integral component of the cortical network involved in processing intraoral thermosensory signals and point to its potential role as a central brain region involved in the integration and communication of behaviorally relevant thermo- and chemosensory information.

## Materials and Methods

### Data acquisition

The experiments in this study were performed on 34 wild type C57BL/6J adult mice (10-20 weeks old; 16 males and 18 females) that were purchased from The Jackson Laboratory (Bar Harbor, ME). Upon arrival to the animal facility, animals were housed on a 12h/12h light-dark cycle and had ad-libitum access to food and water. Experiments and training were performed during the light portion of the cycle. All experiments were reviewed and approved by the Florida State University Institutional Animal Care and Use Committee (IACUC) under protocol “PROTO202100006”.

### Surgery

All animals were anesthetized with an intraperitoneal injection of a cocktail of ketamine (25 mg/kg) and dexmedetomidine (0.25 mg/kg). The depth of anesthesia was monitored regularly via visual inspection of breathing rate, whisker reflexes, and tail reflex. Anesthesia was supplemented by ¼ of the original dose of ketamine as needed throughout the surgery. A heating pad (DC temperature control system, FHC, Bowdoin, ME) was used to maintain body temperature at 35 °C. At the start of surgery, mice were also dosed with dexamethasone (0.4 mg/kg, intramuscular) and bupivacaine HCl (2%, subcutaneous). In addition, lactate solutions were administered every 0.5 h during surgery at volumes of 0.5 ml. Once a surgical plane of anesthesia was achieved, the animal’s head was shaved, cleaned, and disinfected (with iodine solution and 70% alcohol) before being fixed on a stereotaxic holder. To record extracellular activity, mice were implanted with either a custom-made movable bundle of 8 tetrodes [the same used and described in (Vincis et al., 2020; Bouaichi and Vincis, 2020; Neese et al., 2022); n = 28 mice] or with one of two types of chronic and movable silicon probes mounted on a nanodrive shuttle (Cambridge Neurotech). One probe (H5, Cambridge Neurotech; n = 4 mice) had a single shank with 64 electrodes (organized in two adjacent rows spaced 22.5 μm apart) evenly spaced at 25-μm intervals; the other (P1, Cambridge Neurotech; n = 2 mice) had four shanks separated by 250 μm, where each shank had 16 electrodes (organized in two adjacent rows spaced 22.5 μm apart) evenly spaced with 25-μm intervals. Craniotomies were opened above the left GC for implanting tetrodes and probes and above the visual cortex for implanting ground wires (A-M system, Cat. No. 781000). Tetrode bundles, the H5 probes, and the anterior shank of the P1 probes were implanted at AP: +1.2 mm and ML: +3.5 mm (relative to bregma) and were slowly lowered above GC (1.5 mm below the cortical surface). Movable bundles and P1 probes were further lowered 300 μm before the first day of the “control session” recording. H5 probes were further lowered 1200 μm before the first day of the “control session” recording. Tetrodes or probes and a head screw (for the purpose of head restraint) were cemented to the skull with dental acrylic. Before implantation, tetrode wires and the tips of the silicon probes were coated with a lipophilic fluorescent dye (DiI; Sigma-Aldrich), allowing us to visualize tetrode and probe locations at the end of each experiment. Animals were allowed to recover for a minimum of 7 days before the water restriction regimen and training began. Voltage signals from the tetrodes and probes were acquired, digitized, and band-pass filtered with the Plexon OmniPlex system (Plexon, Dallas, TX) (sampling rate: 40 kHz). The time stamps of task events (licking and stimulus delivery) were collected simultaneously through a MATLAB (MathWorks, Natick, MA) based behavioral acquisition system (BPOD, Sanworks) synchronized with the OmniPlex system.

### Experimental Design and Statistical Analysis

#### Behavioral apparatus and training

One week before training began, mice were mildly water restricted (1.5 ml/day) and maintained at or above 85% of their pre-surgical weight. One week after the start of the water restriction regimen, mice were habituated to be head restrained for short (5 minute) daily sessions that gradually progressed (over days) toward longer sessions. During restraint, the body of the mouse was covered with a semicircular opaque plastic shelter to constrain the animal’s body movements without stressful constriction (Fig. 1A). The fluid delivery system, licking detection, and behavioral paradigm have been described in detail in previous studies from our group (Bouaichi and Vincis, 2020; Neese et al., 2022). Following the habituation to restraint, mice were trained with a behavioral task in which the mice learned to lick a dry spout six times to trigger the delivery of the small water drop. Fluid was delivered via gravity by computer-controlled 12 V solenoid valves (Lee Company) calibrated daily to deliver 4 μl from a licking spout (Bouaichi and Vincis, 2020; Neese et al., 2022). A peltier block device (ALA Scientific, NY or custom-made by FSU Machine Shop) located close to the tip of the licking spout was used to heat or cool the water to a specific temperature. The licking spout was made of short polyamide tubing (ID 0.03, MicroLumen) exiting the peltier block. The peltier can be heated or cooled to various temperatures between 0 °C and 50 °C by altering the polarity and the magnitude of DC current provided by a central amplifier. These alterations then led to the heating or cooling of an aluminum block, through which the licking tube passed. To calibrate the temperature of the drinking solution, a thermocouple probe was placed at the exit of the licking spout, and the equipment was considered calibrated when the thermocouple reliably read the desired value of fluid temperature. Therefore, the setting of the temperature (with the associated polarity and value of the DC current) on the central amplifier was based upon the temperature of the fluid exiting the licking spout and not the one within the peltier block. During the habituation and recording sessions (see below for more details), mice received a single 4 μl droplet of deionized water (Barnstead/Thermolyne Nanopure lab water system) at one of three temperatures (14 °C, 25 °C, or 36 °C). These temperatures were chosen for three main reasons: 1) they are outside the range of overt noxious thermal stimuli (Suzuki et al., 2003; Allchorne et al., 2005), allowing the animal to be engaged in the task and actively lick for a substantial number of trials (enabling proper statistical analysis of neural data); 2) they provide compatibility to behavioral studies in rodents that investigated the role of water temperature on intake and preference (Torregrossa et al., 2012; Kay et al., 2020); 3) they represent a broad range of rewarding qualities in water-deprived rodents - with colder stimuli being perceived as more rewarding (Torregrossa et al., 2012). To separate neural activity evoked by the intraoral stimuli from the neural correlates of sensory and motor aspects of licking, we 1) trained the mice to receive each intraoral stimulus, 2) trained the mice for up to two weeks before recording, allowing for familiarization with the different stimuli, and 3) did not analyze any imaging or electrophysiological recording session if the licking pattern evoked by each intraoral stimulus was not similar across at least a 1.5-s temporal window (Fig. 1B), as defined by a Kruskal-Wallis test. As a result, the neural response evoked by the tastants were compared with the response elicited by licking the dry spout before stimulus delivery. This additional requirement also served to ensure that neural activity evoked by the distinct intraoral stimuli would not be impacted by differences in stimulus-evoked licking variables. Water-temperature pairings were presented in a block design, with 10 trials for each block and at least six blocks per session. For the electrophysiology experiments, we relied not only on stereotaxic coordinates and post-hoc evaluation of probe tracks for the successful location of the GC (see Fig. 1C-D) but also on physiological mapping. To this end, we used the “control session” to record the neural activity evoked by different taste qualities and water presented at room temperature (∼ 22 °C). After this recording session, single neuron spiking activity was analyzed. In the case in which taste-responsive neurons were detected, we proceeded with recording GC neurons while mice performed exclusively the “experimental session” for up to three daily sessions. Otherwise the animals were removed from the study (n = 2). At the end of each “experimental session”, tetrodes and P1 probes were further lowered in order to sample new GC neuron ensembles (up to three recording sessions per mouse). For recording sessions with tetrodes, bundles were lowered ∼100 μm after recording each “experimental session”; for recording sessions with P1 probes, they were lowered 200 μm after recording each “experimental session”. For recording sessions with H5 probes, we analyzed only the neural recording obtained during one “experimental session”. In addition to deionized water presented at the selected temperatures mentioned above, the other stimuli used throughout the manuscript include the gustatory stimuli [sucrose (0.1 M), NaCl (0.05 M), citric acid (0.01 M), and quinine (0.001 M)], which were presented at room temperature, and all were purchased from Sigma-Aldrich and dissolved in deionized water to reach the final concentration. For the artificial saliva data, the composition of artificial saliva was based on Breza et al., 2010 (Breza et al., 2010) and consisted of 0.015 M NaCl, 0.022 M KCl, 0.003 M CaCl_2_, and 0.0006 MgCl_2_ with a pH 5.8 ±0.2. Both gustatory stimuli and artificial saliva were freshly prepared each day.

**Figure 1:**
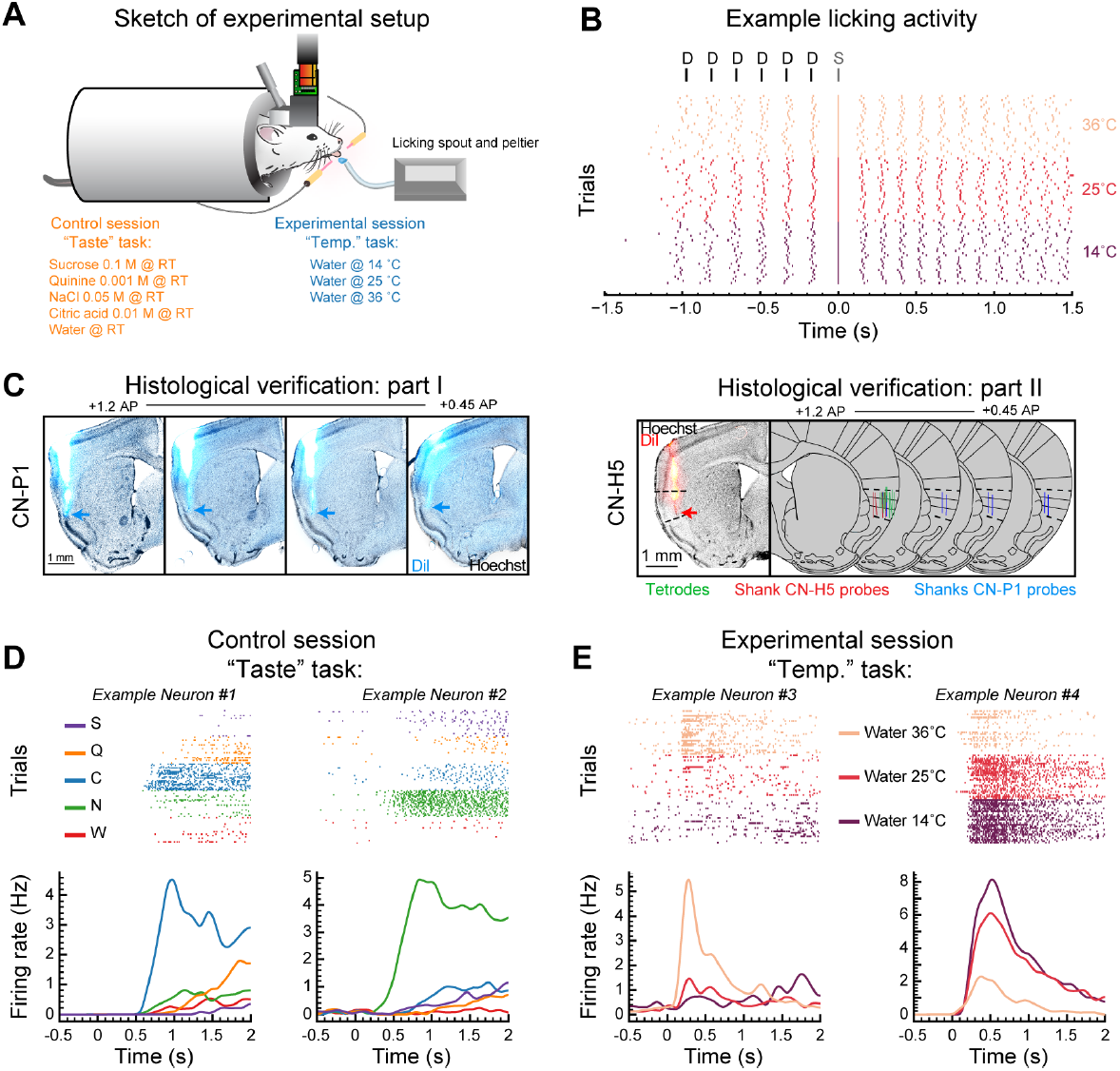
Intraoral thermal responses in the mouse GC. A: Sketch showing the recording setup and a head-restrained mouse licking a spout to obtain intraoral stimuli for two different conditions: the “control” and “experimental” session (see Materials and Methods). B: *top* - diagram of the taste delivery paradigm: intraoral stimuli (S) are delivered after 6 consecutive dry licks (D) to the spout; *bottom* - raster plot of licking activity during one “experimental” session: each line represents an individual lick; trials pertaining to water at different temperature are grouped together and color-coded. Water delivery occurs at time 0 s. C: Example of histological sections showing the tracks (cyan) of the four shanks of the CN-P1 probe in the GC. Cyan arrows point to the tip of the probe. Scale bar is 1 mm. On the right, there is an example of one histological section showing a CN-H5 probe track (red) in the GC. Red arrow points to the tip of the probe. On the far right, there is a schematic of the summary of the tetrode and probe tracks from the 16 mice whose data are analyzed in figures 1-5. Scale bar is 1 mm. D: Raster plots and PSTHs of two representative GC neurons recorded during the “control” session showing taste responses. Trials pertaining to different tastants are grouped together (in the raster plots) and color-coded (both in the raster plots and PSTHs), with sucrose (S) in purple, quinine (Q) in yellow, NaCl (N) in green, citric acid (C) in blue, and water (W) in red. E: Raster plots and PSTHs of two representative GC neurons recorded during the “experimental” session showing intraoral thermal responses. Trials pertaining to water at different temperatures are grouped together (in the raster plots) and color-coded (both in the raster plots and PSTHs).

#### Electrophysiology data and statistical analysis

Kilosort 2 (tetrode data) and Kilosort 3 (probe data) (Pachitariu et al., 2016) were used for automated spike sorting on a workstation with an NVIDIA GPU, CUDA, and MATLAB installed. Following spike sorting, Phy software was used for manual curation. Finally, quality metrics and waveform properties were calculated using code based upon SpikeInterface (Buccino et al., 2020). Only units with an overall firing rate *>* 0.3 Hz, signal to noise ratio *>* 3.0, and an ISI violation rate *<* 0.2 were used for the subsequent analyses. All following analyses were performed on custom Python and R scripts.

##### Water-responsiveness

Water-responsiveness was assessed in all isolated neurons recorded (n = 431 for data presented in figures 1-5; n = 67 for data presented in figure 6; n = 213 for data presented in figures 7-8). This analysis served only to estimate whether and how many GC neurons showed evoked activity that significantly differed from baseline for at least one of the temperatures tested, and not whether there was a significantly different response between the different thermal stimuli. Single-unit spike timestamps were aligned to the stimulus delivery, and peri-stimulus time histograms (PSTHs) were constructed (bin size = 250 ms). Significant changes from baseline were established using a Wilcoxon rank-sum comparison between baseline bin and evoked bin with correction for family-wise error (Bonferroni correction, *p <*0.01). Latency onset was defined as the time in which the smoothed PSTH trace reached half of the max (for active responses where baseline firing rate *<* evoked firing rate) or the min (for suppressed responses where baseline firing rate *>* evoked firing rate) firing rate. For the PSTHs presented in figures 2C and 6C, population responses were obtained by averaging the auROC of each neuron in the observed population. The auROC (area under the receiver-operating characteristic) method normalizes the stimulus-evoked activity to the baseline on a 0–1 scale. A score *>* 0.5 is an active response, and *<* 0.5 is a suppressed response; 0.5 represents the median of equivalence of the baseline activity. For these normalized population PSTHs, a bin size of 100 ms was used.

**Figure 2:**
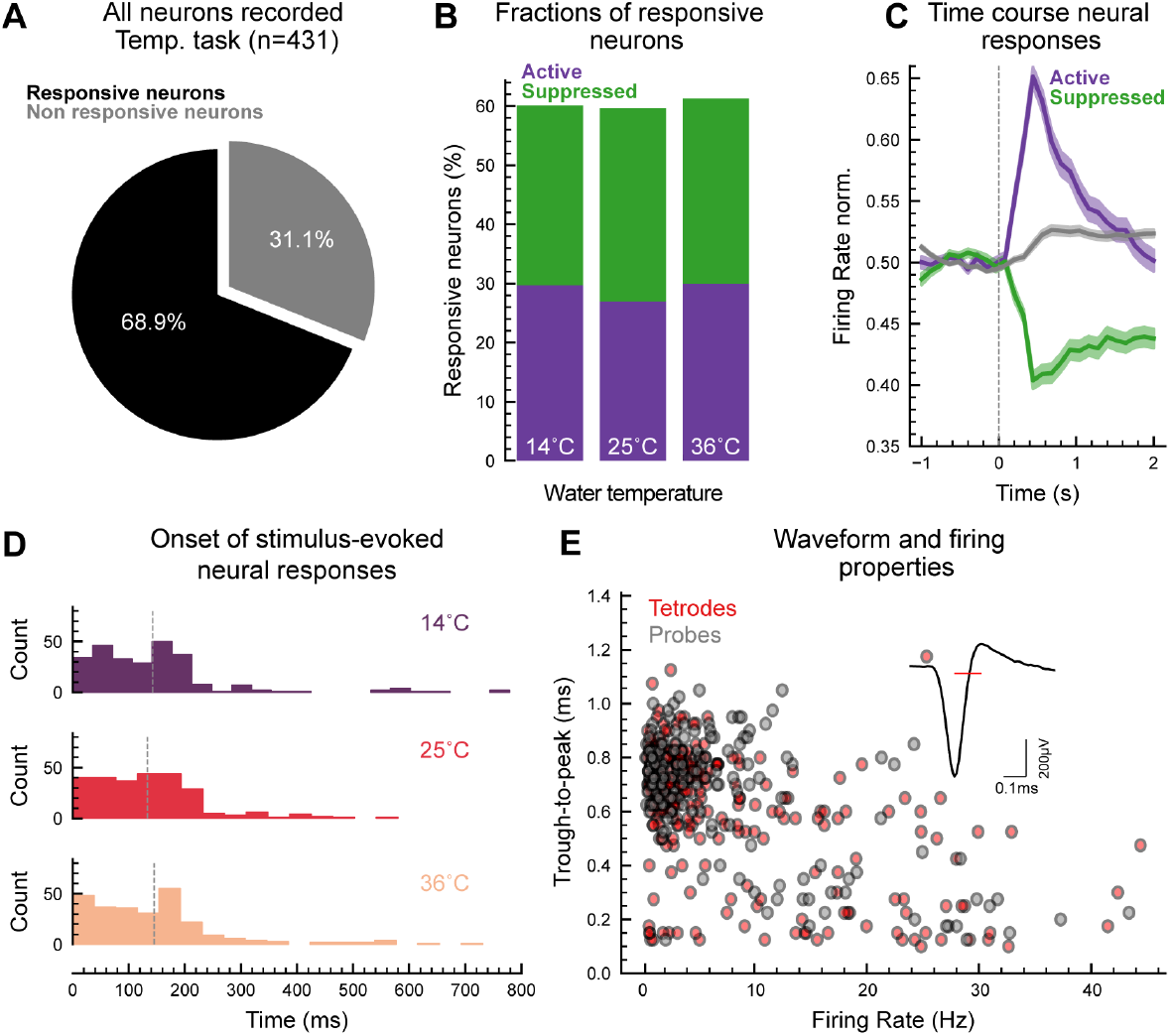
Quantification of water responses in the mouse GC. A: Pie chart showing the proportion of neurons modulated by at least one (14 °C, 25 °C, or 36 °C) intraoral thermal stimulus. B: Bar plots displaying the proportion of responsive neurons showing either an active (*W*.*resp. active* in purple) or suppressed (*W*.*resp. supp*. in green) response to water at 14 °C, 25 °C, or 36 °C. C: Population PSTHs of active (purple), suppressed (green), and nonresponsive (gray) GC neurons expressed as normalized firing rate (norm. FR). Shaded areas represent SE. D: Distribution of onset response latencies. Vertical dashed lines represent mean values. E: Scatter plot of the trough-to-peak duration and firing rate of all single neurons recorded. Red and gray dots represent neurons isolated from tetrode and probe recording sessions, respectively. The inset shows a representative example of a spike waveform with a red horizontal line highlighting the duration of the trough-to-peak interval.

**Figure 3:**
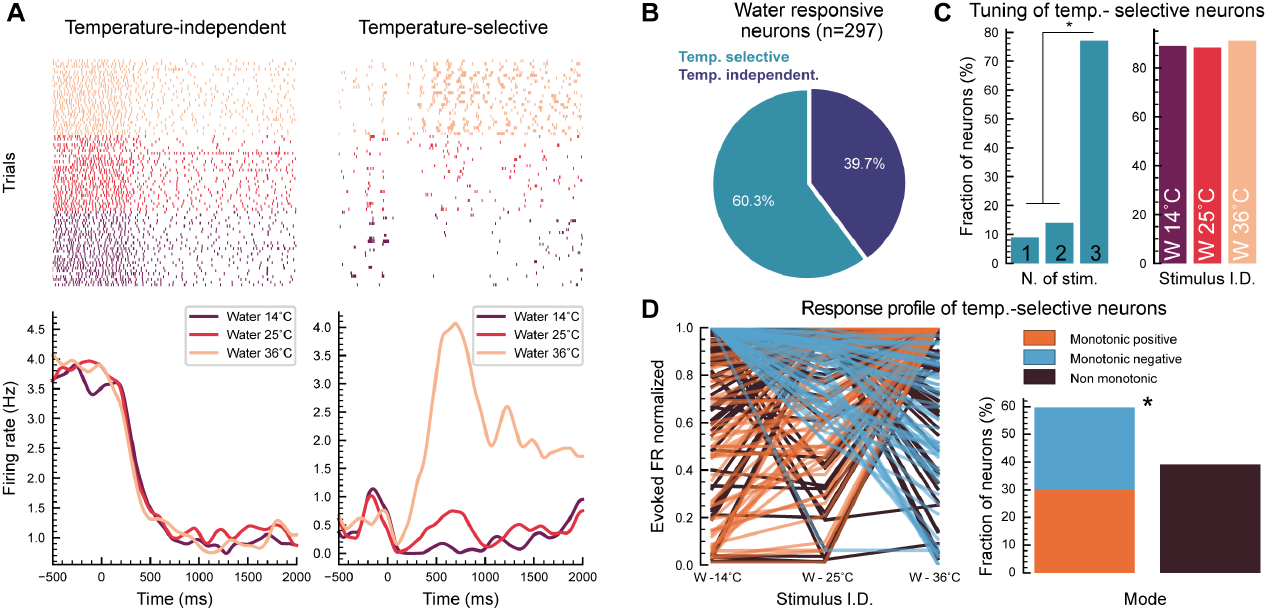
Tuning of temperature-selective neurons in the gustatory cortex (GC). A: Raster plots and PSTHs of two representative water-responsive GC neurons, showing temperature-independent (left) and -selective (right) responses. Trials pertaining to water at different temperatures are grouped together (in the raster plots) and color-coded. B: Pie chart displaying the proportion of water-responsive neurons showing either temperature-selective (in light blue) or -independent (in dark blue) response to water. C: *Left* - Fraction of temperature-selective neurons responding to 1, 2, or 3 thermal stimuli (Chi-squared test for given probabilities: X-squared = 154.94, df = 2, *p*-value *<* 2.2e-16; Multiple proportion comparison - Marascuilo procedure, *p*-value *<* 0.01). *Right* - fraction of temperature-selective neurons responding to the three different temperatures of water (Chi-squared test for given probabilities: X-squared = 0.0875, df = 2, *p*-value = 0.9572). D: *Left* - Plot of peak evoked firing rate (normalized) as a function of water temperature for all the temperature-selective neurons. *Right* - Quantification of the mode (monotonic vs. non-monotonic) of response for temperature-selective neurons in the GC (Chi-squared test for given probabilities: Chi-squared test for given probabilities: X-squared = 7.7345, df = 1, *p*-value = 0.005418).

**Figure 4:**
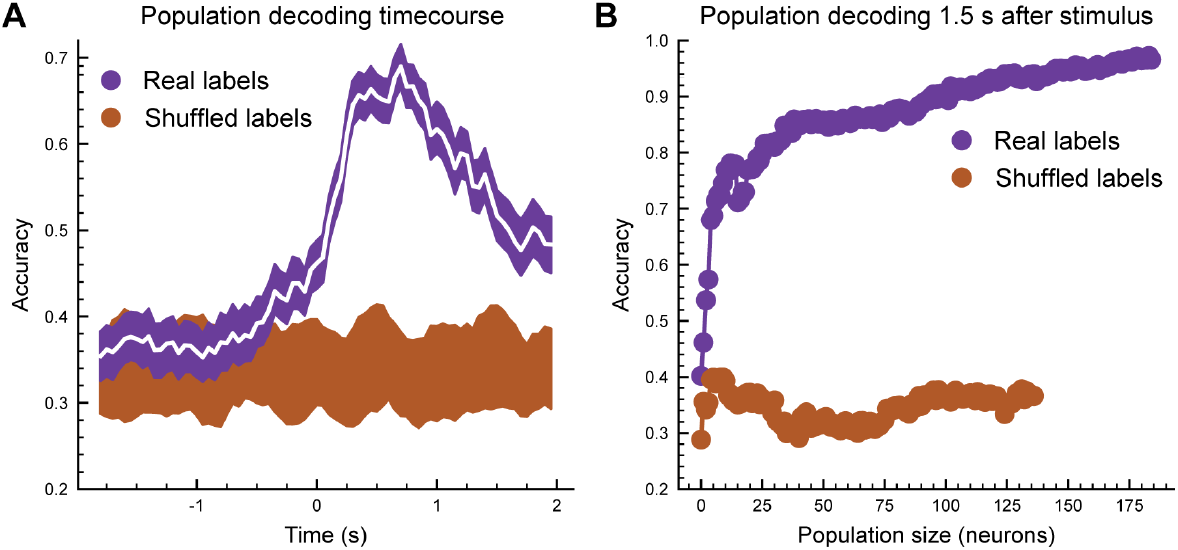
Population decoding of intraoral thermal stimuli in the GC. A: Time course of decoding performance (white line) considering the population of temperature-selective neurons. Purple shaded area indicates the 1%ile-99%ile range of the 20 times the decoder was ran, each time using different training and testing splits (n = 20). Brown shaded areas indicate the 1%ile-99%ile range of the decoding performance over time after shuffling (10 times) stimulus labels for all trials. B: Mean accuracy of SVM linear decoders trained to discriminate the three different temperatures (using a temporal window of 1.5 s after stimulus) as the decoder gained access to progressively more neurons (purple dots). Brown dots indicate the decoding performance over time after shuffling (10 times) stimulus labels for all trials.

**Figure 5:**
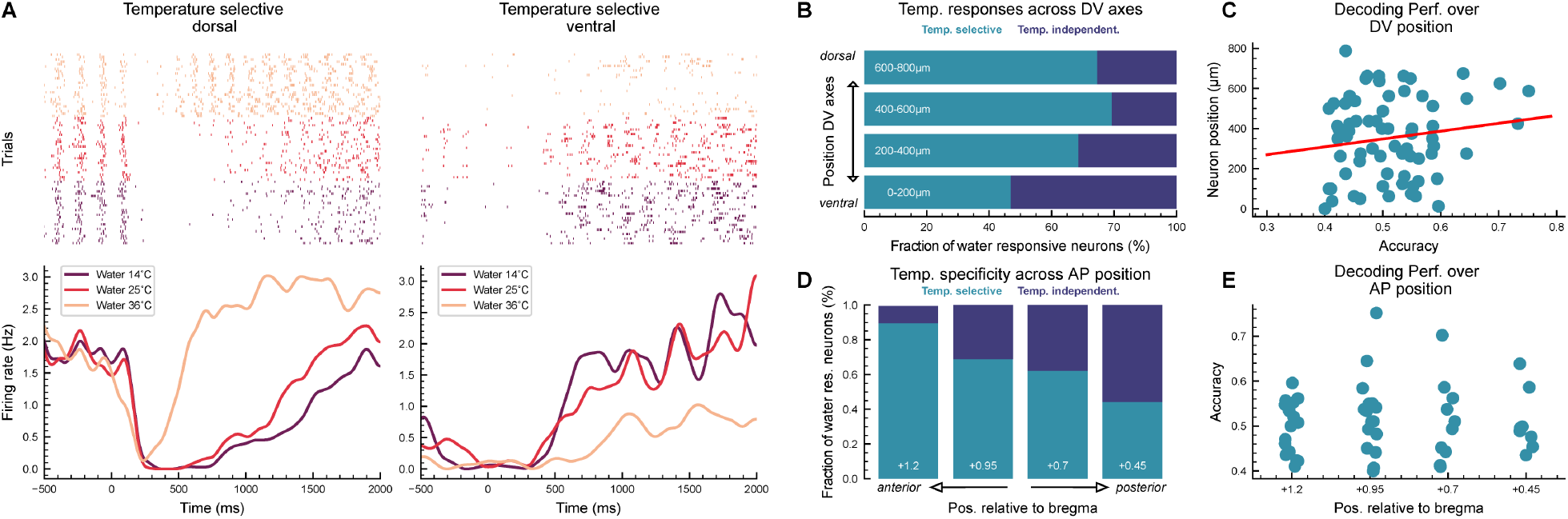
Tuning of temperature-selective neurons across the GC axis. A: Raster plots and PSTHs of two temperature-selective neurons, one recorded in the dorsal (*left*) and one in the ventral (*right*) GC. Water delivery trials at different temperatures are grouped together (in the raster plots) and color-coded (both in the raster plots and PSTHs). B: Fraction of temperature-selective (light blue) and -independent (dark blue) neurons as a function of their dorso-ventral location (four 200-μm bins along the GC anterior-posterior axes; Chi-squared test for given probabilities: X-squared = 8.506, df = 3, *p*-value = 0.03663). C: Scatter plot of the SVM accuracy for temperature-selective neurons against their dorso-ventral location (linear regression analysis: R^2^ = 0.02343, *p*-value = 0.2092). D: Fraction of temperature-selective and -independent neurons as a function of their antero-postero location (Chi-squared test for given probabilities: X-squared = 9.2087, df = 3, *p*-value = 0.02664). E: Plot of the SVM accuracy for temperature-selective neurons against their antero-postero location.

**Figure 6:**
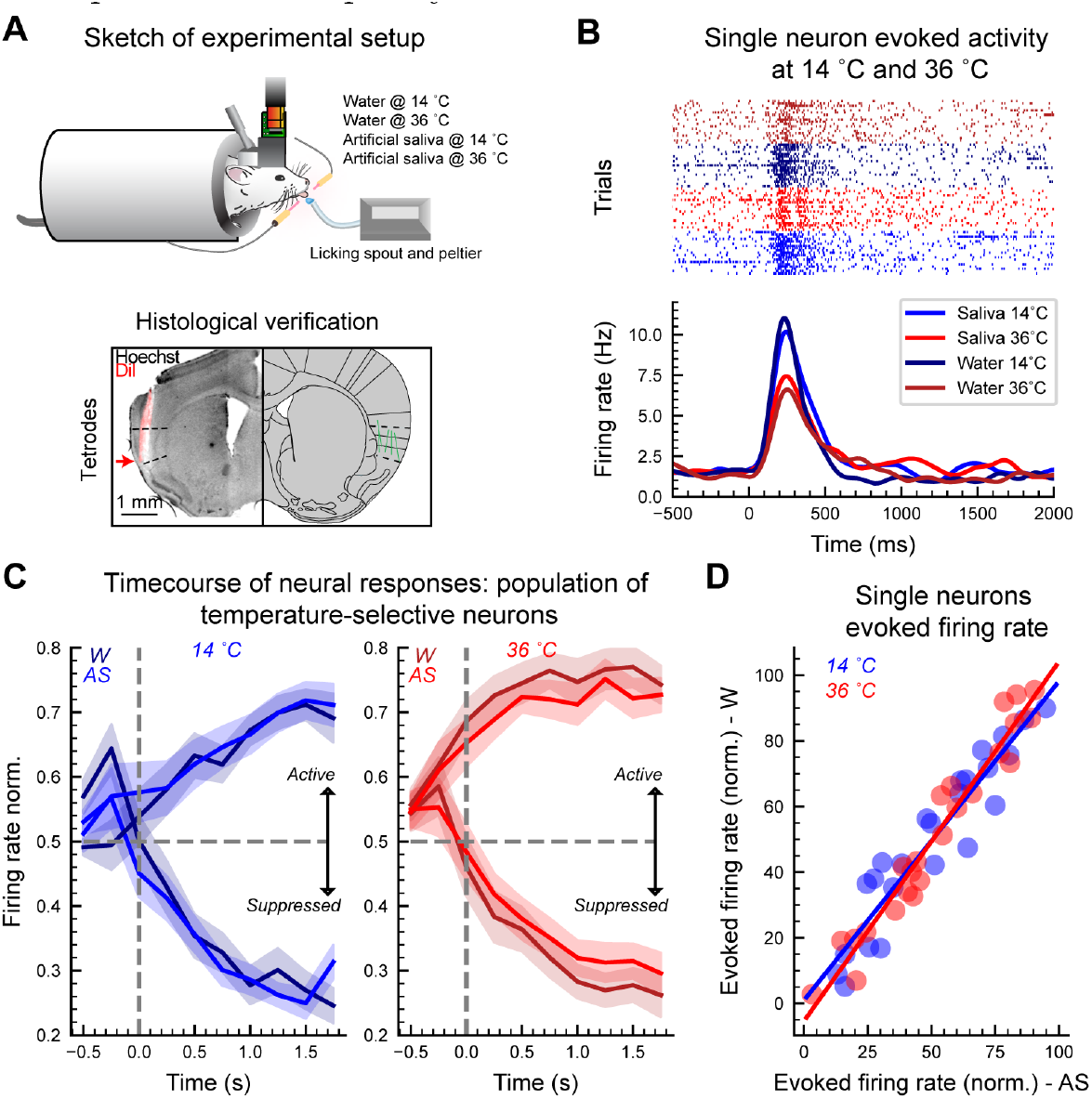
Thermal responses evoked by deionized water and artificial saliva. A: *Top* - Sketch showing the recording setup and a head-restrained mouse licking a spout to obtain deionized water or artificial saliva at two different temperatures (14 °C or 36 °C). *Bottom* - Example of one histological section showing a tetrode track (red) in the GC. Red arrow points to the tip of the tetrodes. On the far right, there is a schematic of the summary of the tetrode tracks from the 5 mice whose data are analyzed in figure 6. B: Raster plot and PSTH of one temperature-selective neuron. Water and artificial saliva delivery trials at different temperatures are grouped together (in the raster plots) and color-coded (both in the raster plots and PSTHs). 36 °C trials are colored with red palette, with darker and lighter red for deionized water (W) and artificial saliva (AS) respectively. 14 °C trials are colored with blue palette, with darker and lighter red for deionized water (W) and artificial saliva (AS), respectively. C: auROC-normalized population PSTHs showing active or suppressed responses after the presentation of deionized water or artificial saliva at 14 °C (*left*) and 36 °C (*right*). Vertical dashed lines indicate stimulus delivery (time = 0 s). Horizontal dashed lines indicate baseline. The shaded area represents the SEM. D: Scatter plot showing the relationship between the firing rate evoked by artificial saliva (x axis) and by deionized water (y axis) at 14 °C (blue) and 36 °C (red). Each dot represents a temperature-selective neuron. The red and blue lines show the linear regression for 36 °C(R^2^ = 0.9481, *p*-value = 7.829e-16) and 14 °C (R^2^ = 0.883, *p*-value = 6.139e-12), respectively.

**Figure 7:**
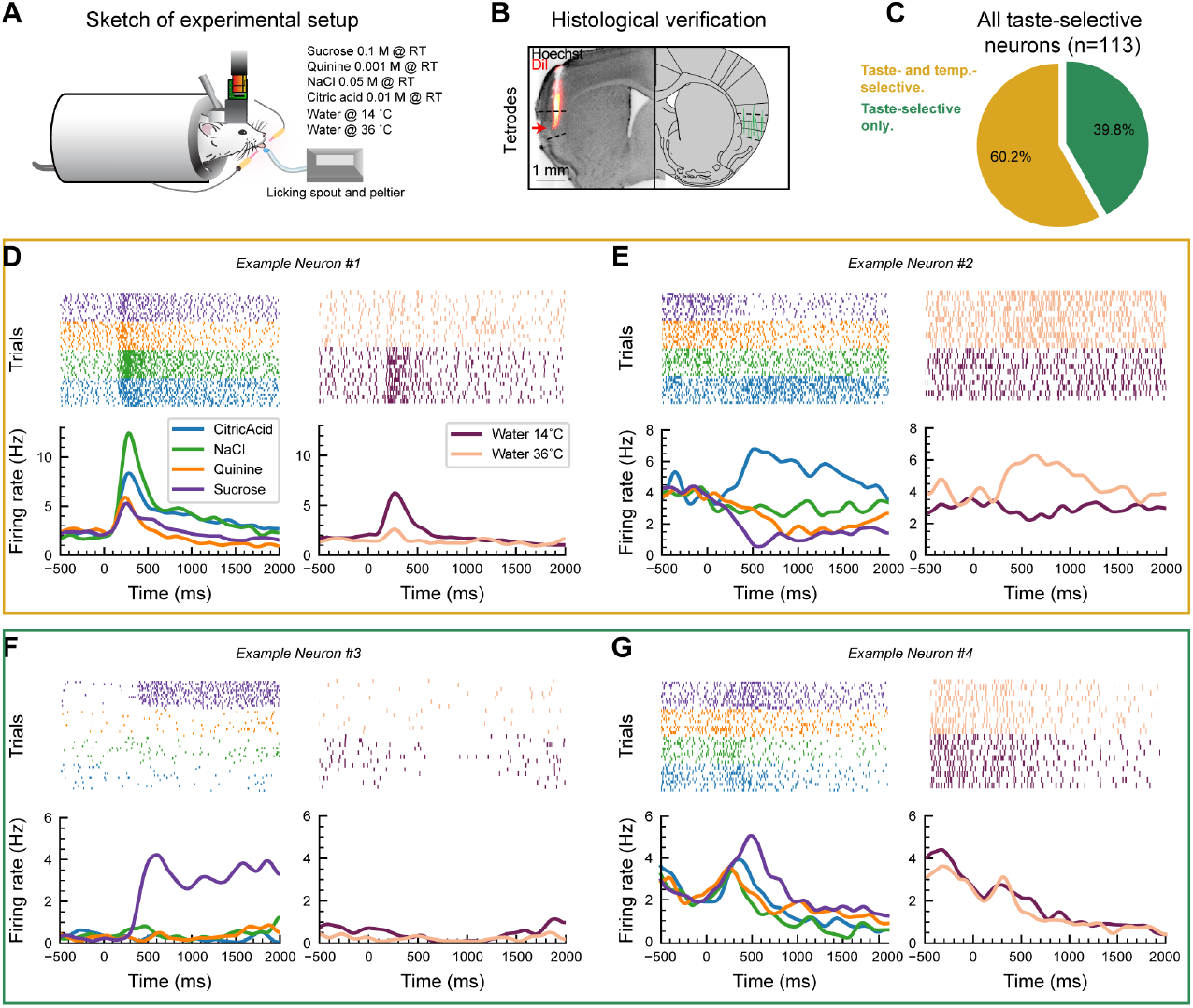
Convergence of chemosensory and thermal intraoral stimuli in the GC. A: Sketch showing the recording setup and a head-restrained mouse licking a spout to obtain deionized water (at 14 °C or 36 °C) or one of four tastants. B: *Left* - Example of one histological section showing a tetrode track (red) in the GC. Red arrow points to the tip of the tetrodes. *Right* - Schematic of the summary of the tetrode tracks from the 12 mice whose data are analyzed in figure 7. C: Pie chart showing the proportion of taste-selective neurons (n = 113) that are modulated exclusively by taste (taste-selective only) or by taste and thermal stimuli (taste- and temperature-selective). D-E: Raster plots and PSTHs of two taste- and temperature-selective neurons (one neuron in panel D and one neuron panel E). Taste (*leftmost*) and water (*rightmost*) trials are separated for clarity. F-G: Raster plots and PSTHs of two taste-selective only neurons (one neuron in panel F and one neuron panel G). Similar to panels D-E, taste (*leftmost*) and water (*rightmost*) trials are separated for clarity.

**Figure 8:**
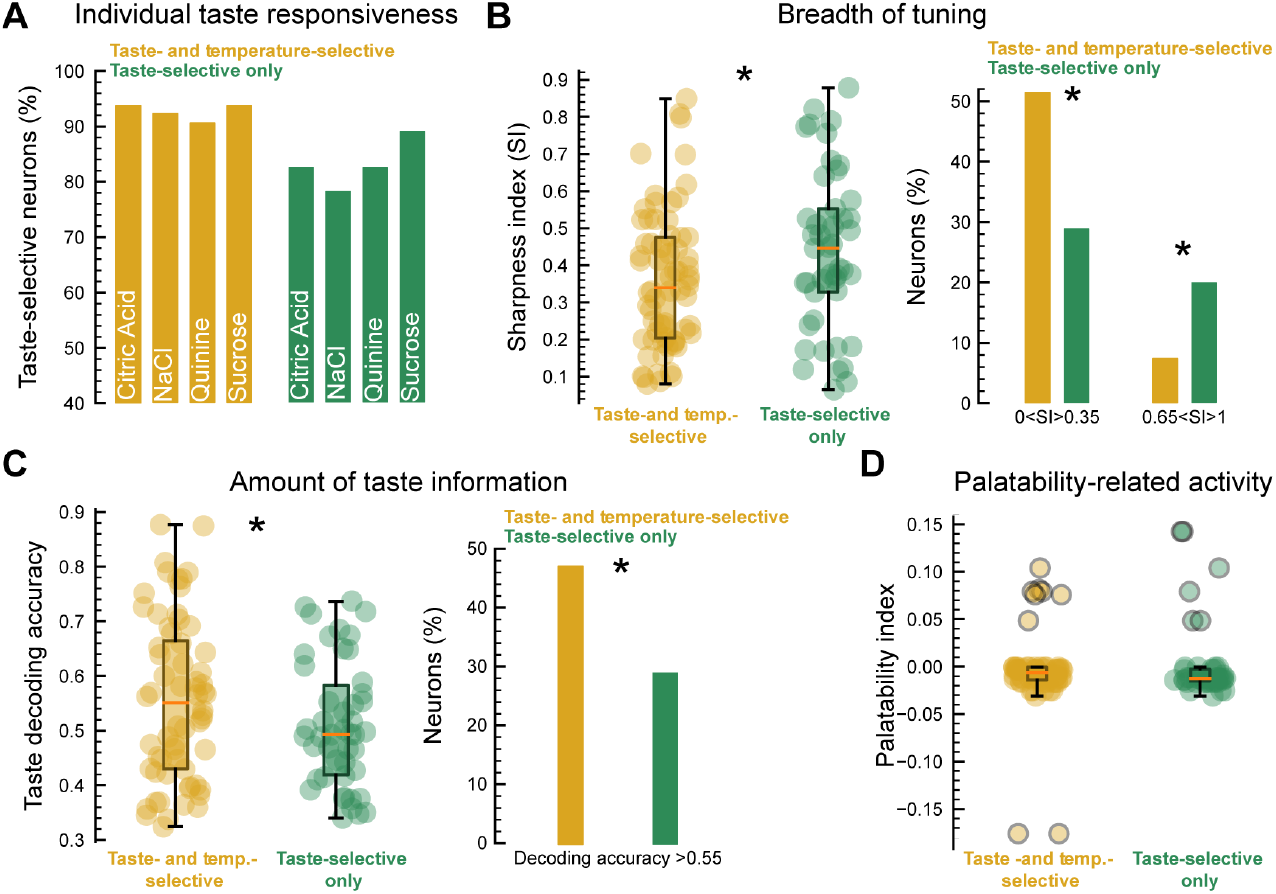
Taste-selective properties of the two subsets of GC neurons. A: Fraction of taste- and temperature-selective (gold) and taste-selective only (green) neurons responding to the four different taste qualities. B: *left* - Box plots showing the distribution of the breadth of tuning (expressed as sharpness index, SI) of the taste- and temperature-selective (gold) and taste-selective only (green) neurons. High SI values indicate narrowly responsive neurons, whereas low SI values imply that the neuron is modulated by multiple tastants. Each colored circle represent a single neuron. *right* - Fraction of taste- and temperature-selective (gold) and taste-selective only (green) neurons with a low (0*<*SI*>*0.35) and high (0.65*<*SI*>*1) SI values range (low SI range, 2-sample test for equality of proportions: X-squared = 4.137, df = 1, *p*-value = 0.02098; high SI range, 2-sample test for equality of proportions: X-squared = 2.9103, df = 1, *p*-value = 0.04401). C: *left* - Box plots showing the distribution of the taste decoding accuracy of the taste- and temperature-selective (gold) and taste-selective only (green) neurons. Each colored circle represent a single neuron. *right* - Fraction of taste- and temperature-selective (gold) and taste-selective only (green) neurons with a high (*>* [chance-level ×2]) decoding accuracy for taste information (2-sample test for equality of proportions: X-squared = 3.011, df = 1, *p*-value = 0.04135). D: Box plots showing the distribution of the palatability-related firing rate (expressed as palatability index, PI) of the taste- and temperature-selective (gold) and taste-selective only (green) neurons. Each colored circle represents a single neuron. Circles outlined in black represent outliers.

##### Temperature-selective neurons

To determine the degree of temperature specificity of GC water responses, we subjected each of the water-responsive neurons to a support vector machine (SVM) classifier, used and described in detail in a previous study (Neese et al., 2022).

Briefly, the spike train dataset (0-1.5 s after water delivery) was transformed into a collection of vectors, each corresponding to one experimental trial. Each trial in the dataset was classified hierarchically according to neuron ID and temperature of water (labels: 14 °C, 25 °C, or 36 °C) for the trial. Thus, several vectors were associated with a single neuron, depending on how many trials were run with different temperatures. Then, for each single neuron, the classification analysis consisted of separating the ensemble of vectors into a training set consisting of 67% of the spike trains, and a testing set consisting of the remaining 33% of the spike trains. The training set was used to fit parameters for the SVM model (linear SVM kernel was used throughout our analysis); the SVM searched for hyperplanes that best separated the various classes (i.e., points corresponding to trials for different water temperatures) within the dataset. The trained model was then used to classify the testing dataset, resulting in a classification score measured as the percentage of correctly classified points. This procedure was repeated 20 times for each neuron, each time using a different partition of vectors into training and testing sets, and the classification scores were averaged over these trials to obtain an overall classification score for the neuron. To assess the significance of the overall classification score, we used a permutation test where the labels of the trials were shuffled without replacement. Spike trains were shuffled 100 times, and the pseudo classification score index was calculated for each iteration of the shuffling. A neuron was deemed temperature-selective if its overall classification score was *>* 99%ile of the pseudo classification scores (*p <*0.01).

##### Population decoding classifier

To understand how well the GC encoded information regarding intraoral temperature (Fig. 4), we used a population decoding approach. To this end, we first constructed a pseudopopulation of GC neurons using temperature-selective neurons recorded across different sessions (n = 179). We then generated a firing rate matrix (trials × time bin) where the spike timestamps of each neuron (2 s before and 2 s after stimulus) were realigned to water delivery, binned into 50-ms time bins. To assess the amount of temperature-related information, we used the SVM classifier described above. Spike activity data contained in our matrix were divided into a training set consisting of 67% of the spike trains and a testing set consisting of the remaining 33% of the spike trains. This process was repeated 20 times (each time using different training and testing splits) to compute the decoding accuracy, defined as the fraction of trials in which the classifier made correct temperature predictions using a 25-ms sliding window. To assess the significance of the population decoding over time, we used a permutation test where the labels of the trials were shuffled without replacement. Spike trains were shuffled 10 times, and the pseudo classification score over time was calculated for each iteration of the shuffling.

##### Taste-selective neurons

In order to define a neuron as taste-selective, two criteria must be satisfied: (1) activity significantly differs from baseline and (2) there is a significantly different response among the four tastants. Significant changes from baseline were evaluated using Wilcoxon rank-sum comparison as already described in the *Water-responsiveness* sub-section of the Method section. Significant changes between tastes were determined using a support vector machine (SVM) classifier as already described in the *Temperature-selective neurons* sub-section of the Method. To further investigate the taste response profile of taste-selective neurons, we used sharpness (SI) (Rainer et al., 1998; Yoshida and Katz, 2011; Bouaichi and Vincis, 2020) and palatability (PI) (Piette et al., 2012; Bouaichi and Vincis, 2020) indices. SI was computed on the mean firing rate during the 1.5-s-wide interval after taste delivery and was defined as

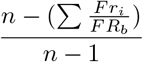

where *Fr*_*i*_ is the mean firing rate for each taste (*i* = 1–4), *FR*_*b*_ is the maximum firing rate among gustatory stimuli, and *n* is the total number of stimuli (*n* = 4). A SI of 1 indicated that a neuron responded to one stimulus (narrow tuning), and the value 0 indicated equal responses across stimuli (broad tuning). To evaluate whether taste-selective neurons encoded palatability-related information, we used the palatability index (PI). To avoid potential confounds introduced by differences in baseline and evoked firing rates across our pools of taste-selective neurons, we first normalized the PSTHs with the auROC procedure. We then computed the absolute value of the log-likelihood ratio of the normalized firing rate for taste responses with similar (|LR|_*same*_) and opposite (|LR|_*opposite*_) hedonic values:

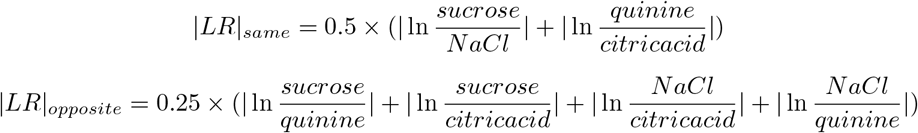

We then defined the PI as |LR| _*opposite*_-|LR| _*same*_. Positive PI values suggested that a neuron responded similarly to tastants with similar palatability and differently to stimuli with opposite hedonic values. Negative PI values indicated the alternative scenario in which a neuron responded differently to stimuli of the same palatability and similarly to taste with different hedonic values.

#### Histology

At the end of the experiment, mice were terminally anesthetized and perfused transcardially with 30 ml of PBS followed by 30 ml of 4% paraformaldehyde (PFA). The brains were extracted and post-fixed with PFA for 24 h, after which coronal brain slices (100-μm thick) containing the GC were sectioned with a vibratome (VT1000 S; Leica). To visualize the tracks of the tetrode bundles and probes, brain slices were counterstained with Hoechst 33342 (1:5,000 dilution, H3570; ThermoFisher, Waltham, MA) by standard techniques and mounted on glass slides. GC sections were viewed and imaged on a fluorescence microscope.

## Results

### Single neuron analysis of GC intraoral thermal responses

To investigate how single GC neurons encode intraoral thermal information in freely licking mice, we recorded ensembles of single units via movable bundles of tetrodes (Vincis et al., 2020; Bouaichi and Vincis, 2020; Neese et al., 2022) or movable silicon probes (Cambridge Neurotech) mounted on a nanodrive shuttle (Cambridge Neurotech) implanted unilaterally in the GC (Fig. 1A). After habituation to head restraint, water-deprived mice were engaged in a behavioral task in which they had to lick 6 times to a dry spout to obtain a 4-μl drop of one intraoral fluid stimulus (Fig. 1B). On alternating days for up to 10 days, mice were trained to receive either five gustatory stimuli (0.1 M sucrose, 0.05 M NaCl, 0.01 M citric acid, 0.001 M quinine, and water) at room temperature (“control session”; Fig. 1A) or only water at three different temperatures (14 °C, 25 °C, or 36 °C; “experimental session”; Fig. 1A). During this training, tetrode bundles or silicon probes were slowly lowered to their final position (Fig. 1C) as described in the Methods section. At the end of the training, the recording session began. First, we recorded neural activity evoked by different taste qualities and water presented at room temperature (“control session”; Fig. 1D). After this recording session, single neuron spiking activity was analyzed; if taste-selective neurons were detected, in the next recording sessions, we exclusively recorded activity evoked by water at the different temperatures (“experimental session”; Fig. 1E; 3 recording sessions for all mice except the ones implanted with the H5 probe; at the end of each experimental session, tetrodes and probes were lowered to record new ensembles of neurons [see Materials and Methods section for more details]). It is important to highlight we are not claiming to have tracked the same taste-selective neurons across multiple days and recording sessions. Rather, this approach was chosen exclusively to provide additional functional evidence to support that recordings aimed at neuron responses to orally-sourced thermal stimuli (“experimental session”; Fig 1E) were indeed obtained from the GC. A total of 431 single neurons were recorded from 16 mice with an average yield of 11.42 ±6.49 neurons per session.

To begin evaluating the neural dynamics evoked by intraoral thermal stimuli in active licking mice, we analyzed the spiking profile of single GC neurons. Figure 1E shows the raster plots and PSTHs of two representative GC neurons. Visual inspection of the graphs indicated that each of these neurons was modulated by different temperatures of water (Fig. 1E). As a first step, we wanted to understand how many GC neurons were modulated by the presence of a solution in the mouth. This analysis was performed by comparing the baseline and evoked neural activity from each of the three water temperatures and therefore serves only to estimate whether and how many GC neurons were responsive to intraoral thermal stimuli—not whether those responses differ as a function of the temperature of the stimulus. Wilcoxon rank-sum analysis revealed that a substantial number of the recorded GC neurons [68.9% (297/431)] responded to at least one of the three intraoral thermal stimuli and were classified as “water-responsive” (Fig. 2A). We observed a similar fraction of water-responsive neurons showing active and suppressed activity to each of the temperatures tested (Fig. 2B). Figure 2C shows the population averages (population PSTHs) of the active and suppressed responses. When comparing the sign of water-responsive neurons, we observed significant differences between the number of water-responsive neurons that show an active vs. suppressed response.

Analysis of the distribution of the latency of the responses indicated three main points. First, the majority of water-responsive neurons showed a fast onset, with firing rate significantly changing from baseline within the first 200 ms after stimulus delivery (Fig. 2D; mean onset 0.29 ±0.360 s). Second, the distribution of response latencies does not differ as a function of the temperature of the stimulus (Kruskal-Wallis chi-squared = 0.087628, df = 2, *p*-value = 0.9571). Third, analysis of the distribution of two waveform properties (firing rate and trough-to-peak) revealed no differences in the population of neurons sampled via tetrodes and silicon probes (Fig. 2E; firing rate comparison, t-test: t = 0.92692, df = 380.99, *p*-value = 0.3546; trough-to-peak comparison, t-test: t = -1.7226, df = 427.94, *p*-value = 0.08569).

Next, we wanted to investigate the relationship between neural activity and oral thermosensation. To this end, we evaluated whether the evoked activity of water-responsive neurons differs as a function of the temperature of the water. A qualitative evaluation of the raster plots and PSTHs in figures 1 and 3 indicated that these responses can be classified as temperature-independent (i.e., water responses appear to be similar at the three temperatures tested; see for example Fig. 3A *left*) and as temperature-selective (i.e., evoked responses appear to differ as a function of the thermal stimulus; see for example Fig. 1E and Fig. 3A-*right*). To quantify the degree of temperature selectivity of GC neurons, we subjected each of the water-responsive neurons to a support vector machine (SVM) classifier (Neese et al., 2022; see Materials and Methods for more details). The SVM uses a supervised machine learning algorithm that acts as a classifier whose performance was used as a surrogate for the ability of each individual neuron to encode thermal information in a 1.5-s post-stimulus temporal window. This analysis revealed that more than half of the GC water-responsive neurons (60.3%, 179/297) were temperature-selective (Fig. 3B), whereas the remaining (39.7%, 118/297) likely encoded for thermal-independent features. We then wanted to understand if the temperature-selective GC neurons were preferentially modulated by only a specific temperature or if they were capable of encoding information pertaining to multiple thermal stimuli. This analysis serves only to estimate how many temperature-selective neurons were modulated by more than one thermal stimulus, independent of the degree of the temperature. The distribution plot shown in the left panel of figure 3C indicates that the majority of temperature-selective neurons were capable of responding to all three of the temperatures tested (Fig. 3C-*left* ; Chi-squared test for given probabilities: X-squared = 154.94, df = 2, *p*-value *<* 2.2e-16; Multiple proportion comparison - Marascuilo procedure, *p*-value *<* 0.01).

Next, we wanted to evaluate if GC temperature-selective neurons were tuned to both cooling and warming intraoral stimuli or if they preferentially responded to either one. When we considered the absolute values of the water stimulus (14 °C, 25 °C, or 36 °C), we observed that temperature-selective neurons were broadly responsive to all thermal stimuli, independent of the degree (°C) value (Fig. 3C-*right* ; Chi-squared test for given probabilities: X-squared = 0.0875, df = 2, *p*-value = 0.9572). If instead we consider deviation from resting oral temperature (32 °C (Leijon et al., 2019); Δ*T*), more GC neurons appear to be tuned to cooling (*<*32 °C) than to warming (*>*32 °C) stimuli. However, this latter observation might be biased due to the limited and uneven set of stimuli used (14 °C and 25 °C are both “cooling” and 36 °C is “warming”; see the Discussion section for more on this point). To further investigate how GC temperature-selective neurons were tuned to absolute changes in orally-sourced temperature of fluid solution, we performed an additional analysis: we evaluated if the neuron’s firing either positively or negatively correlated with increase of temperature (monotonic) or not (non-monotonic). Our analysis revealed that the majority of temperature-selective neurons changed their firing rate in a monotonic fashion (Fig. 3D; Chi-squared test for given probabilities: X-squared = 7.7345, df = 1, *p*-value = 0.005418). Overall, these analyses revealed that most of the GC neurons that responded to the delivery of water were temperature-selective and capable of encoding different absolute temperature values in a mostly monotonic manner.

### Population coding for intraoral temperature in GC

To further characterize cortical thermosensory processing, we performed a population decoding analysis. The decoder was instantiated using the same SVM classifier described above for single neurons; the difference in this case is that the classifier was trained using single trial responses of pseudopopulations of temperature-selective GC neurons (neurons pooled from different experimental sessions and animals) and tested using a held-out method (training set consisting of 67% of the spike trains and a testing set consisting of the remaining 33% of the spike trains). This process was repeated 20 times (each time using different training and testing splits) to compute the decoding accuracy, defined as the fraction of trials in which the classifier made correct temperature predictions. We began by analyzing how well the population activity of temperature-selective neurons (n = 179) represented the three thermal stimuli over a 2-s long post-stimulus temporal window (Fig. 4A). Figure 4A shows the decoding performance of the pseudopopulation using a sliding window of 25 ms (white trace [average accuracy over the 20 training and testing splits] over the purple shading [1%ile (lower bound) and 99%ile (upper bound) of the 20 training and testing splits]). As a control, the same analysis was performed another 10 times after shuffling the thermal stimuli labels; the brown shaded area in Figure 4A shows the 1%ile (lower bound) and 99%ile (upper bound) of the distribution of the control decoding values over time. The decoding time-course showed an early onset (classification above control) and reached its peak within 1 s after stimulus delivery. Additionally, although the overall classification value started decreasing after 500 ms, decoding performances remained above control until the end of the temporal window analyzed (Fig. 4A). Figure 4B shows how classification during a 1.5-s post-stimulus temporal window changed as the decoder gained access to progressively more neurons. While the classifiers that trained with a small numbers of neurons were less accurate at identifying intraoral thermal information, increasing the number of neurons in the pseudopopulation drastically increased the decoding performance, reaching up to 85% accuracy with less than 40 neurons (Fig. 4B, purple dots). As expected, the decoding performance obtained with the control data did not improve and stayed around chance level even when the decoder gained access to progressively more neurons (Fig. 4B, brown dots). Taken together, these results indicate that both individual and ensembles of temperature-selective GC neurons rapidly and reliably encoded intraoral thermosensory signals over time, with classification accuracy for stimulus identity remaining above chance and control level for up to 2 s after stimulus delivery.

### Spatial organization of intraoral thermal responses

Next, we sought to determine whether intraoral responses were topographically organized. A previous study in anesthetized rats suggested that thermal responses in the GC are mostly clustered in the dorsal (granular) region (Kosar et al., 1986). However, it is still unknown whether intraoral thermal responses in behaving mice are also organized with a seemingly topographical gradient along the dorso-ventral axes of the GC.

To address this question, we took advantage of the recording sessions performed with chronic probes that allowed us to triangulate the location of the recorded neurons along the dorso-ventral axes spanning up to 800 μm (see Materials and Methods section for more details). Visual inspection of spiking activity shown in Figure 5A indicated the presence of temperature-selective neurons in the dorsal and ventral region of the GC. We divided the dorso-ventral axes of the portion of the GC captured by our recording into four 200-μm spatial bins (Fig. 5B-C). Our post-hoc evaluation of the implant tracks confirmed that the most ventral position was well within the agranular region of the GC (Fig. 2B). Overall, our probe recordings spanned 800 μm, which covered a large portion of the dorso-ventral plane of the mouse GC (Wang et al., 2020). Analysis of the temperature-selective neurons along the GC dorso-ventral axes reflected a coarse topographical organization with much fewer temperature-selective neurons in the most ventral part of GC (0-200 μm) compared to the other three spatial bins analyzed (Fig. 5B; 4-sample test for equality of proportions: X-squared = 8.506, df = 3, p-value = 0.03663). To further investigate between temperature-selective neurons and their location along the GC dorso-ventral axes, we plotted the single neuron SVM decoding accuracy against the neuron position (Fig. 5C). We reasoned that a topographical clustering might also appear when considering the amount of thermal-related information (expressed as decoding accuracy) of the temperature-selective neurons (for example, neurons with high accuracy could cluster more dorsally). Contrary to our prediction, visual inspection of Figure 5C and linear regression analysis (R^2^ = 0.02343, *p*-value = 0.2092) revealed that was not the case. We then focused our attention on the organization of intraoral thermal responses along the rostro-caudal axes. Recent studies have started to uncover the role of the mouse posterior insular cortex in skin warming and cooling (Vestergaard et al., 2023; Beukema et al., 2018). To determine if intraoral thermal responses in the GC are organized in a antero-postero gradient, we analyzed the temperature-selective neurons recorded with the P1 probe (Fig. 1D) that allowed us to examine responsiveness along 1 mm of the GC antero-postero axes. Analysis of the distribution of temperature-selective neurons (Fig. 5D) revealed an antero-postero gradient with more temperature-selective neurons in the anterior GC (Fig. 5D; 4-sample test for equality of proportions: X-squared = 9.2087, df = 3, *p*-value = 0.02664). However, similar to what we observed for the dorso-ventral axes, the capability of GC neurons to decode thermal information did not depend on their position along the antero-postero axes (Fig. 5E ; one way ANOVA: F(3,48) = 0.239, *p*-value = 0.869).

In summary, these results suggest that the number of temperature-selective responses are, for the most part, sensitive to anatomic location within the mouse GC. The ratio of temperature-selective and temperature-independent neurons indicate there is a coarse organization along the dorso-ventral and antero-postero axes. However, taste-sensitive neurons appear to encode thermal information (as revealed by the SVM classifier) equally well no matter location within the mouse GC.

### Comparison of intraoral thermal activity evoked by deionized water and artificial saliva

Water-specific responses have been reported in many brain regions of the gustatory neuraxis (Zocchi et al., 2017; Nakamura and Norgren, 1991; Gutierrez et al., 2010; Bouaichi and Vincis, 2020; Neese et al., 2022; Rosen et al., 2010); in addition, previous studies have argued (Accolla et al., 2007) and provided evidence (Zocchi et al., 2017) in favor of water as an independent taste quality.

Therefore, we sought to understand if the temperature-selective activity observed so far in the GC neurons was mostly driven by thermal information in absence of an overt taste stimulus (as opposed to “taste”-temperature integration). To this end we designed an experiment that allowed us to compare thermal responses with water to those with artificial saliva - another intraoral stimulus often used as a neutral control instead of deionized water in taste research (Breza et al., 2010; Baumer-Harrison et al., 2020; Travers et al., 2022). We recorded neural activity from GC neurons (n = 67) of a second group of mice (n = 5) trained to receive deionized water or artificial saliva at 14 °C or 36 °C (Fig. 6A). Figure 6B shows the raster plots and PSTHs of one temperature-selective GC neuron. A qualitative analysis of the single neuron’s thermal responses indicated that neural activity evoked by deionized water and artificial saliva was very similar (Fig. 6B). Next, we wanted to evaluate the similarity between the activity evoked by water and artificial saliva at the two temperatures in all temperature-selective neurons. Figure 6C shows the auROC-normalized population (all temperature-selective neurons) averages of the time-course of the responses to 14 °C and 36 °C, further highlighting the similarity of the neural activity evoked by deionized water and artificial saliva (Fig. 6C). To perform a quantitative evaluation of the similarity of the neural activity, we used a linear regression to test if the evoked firing rate in the 1.5-s temporal window - with water as stimulus - significantly predicted the evoked firing rate of artificial saliva (Fig. 6D). For both 14 °C and 36 °C, the overall regression was statistically significant (14 °C: R^2^ = 0.883, *p*-value = 6.139e-12; 36 °C: R^2^ = 0.9481, *p*-value = 7.829e-16), indicating that temperature-selective neurons in the GC appear to encode intraoral thermal information in absence of overt chemosensory taste stimuli.

### Evaluating the convergence of thermal and chemosensory information on GC neurons

While our results thus far have described how orally-sourced thermal inputs are processed in the GC, this cortical region is typically studied for its role in processing chemosensory taste information (Katz et al., 2001; Jezzini et al., 2013; Bouaichi and Vincis, 2020; Neese et al., 2022; Stapleton et al., 2006). In addition, recent electrophysiological studies suggest that GC neurons are capable of integrating multimodal intraoral signals (Vincis and Fontanini, 2016a; Samuelsen and Vincis, 2021; Maier, 2017).

Therefore, we next explored whether GC neurons responsive to thermal variation represent a unique population of cells embedded among taste-selective cells or whether - and to what extent - thermal and chemosensory information converge on the same neurons. To accomplish this, we recorded neural activity from GC neurons of a third group of mice (n = 12; 4 of these mice were also used for data presented in figure 6) trained to receive - in the same experimental session - a palette of six intraoral stimuli: four chemosensory taste stimuli presented at room temperature (sucrose, 0.1 M; NaCl, 0.05 M; citric acid, 0.01 M; quinine, 0.001 M) and two thermal stimuli (deionized water at 14 °C and 36 °C) (Fig. 7A-B). The concentration and temperature of the taste stimuli were chosen to provide compatibility to prior awake-behaving taste electrophysiology studies in mice (Bouaichi and Vincis, 2020; Neese et al., 2022; Levitan et al., 2019; Dikecligil et al., 2020). Out of the 213 single neurons recorded, 113 were taste-selective. Taste-selective neurons were required to satisfy two criteria: they must have exhibited a significant firing rate change from baseline (defined by Wilcoxon rank-sum), and they must have shown significantly different responses to the four tastants (defined by the SVM analysis). Further, more than half (60.2%, 68/113) of the taste-selective neurons were also temperature-selective (Fig. 7C); these neurons will hereafter be referred to as taste- and temperature-selective neurons. Figure 7D-E shows the raster plots and PSTHs of two representative taste- and temperature-selective neurons. Visual inspection of the graphs indicated that each of these neurons was modulated by different taste stimuli and also by the two different temperatures of water. On the contrary, 39.8% (45/113) of taste-selective neurons were not modulated by the thermal inputs (Fig. 7C) and were therefore been deemed taste-selective only neurons. Representative examples of this category of GC neurons are shown in figure 7F-G. These qualitative results started to paint a picture wherein the majority of GC neurons that encoded taste information were also capable of integrating intraoral thermal stimuli. To further explore this idea, we performed a series of analyses aimed at understanding whether the GC neurons that responded to both taste and thermal information represented a unique population of cells embedded among taste-selective cells. First, we aimed to evaluate if either of the two sets of GC neurons (taste- and temperature-selective neurons and taste-selective only neurons) were differently tuned to encode information pertaining to individual taste qualities. Figure 8A shows that both groups of neurons responded to gustatory stimuli independent of their chemical identity, and the total fraction of taste-selective only neurons modulated by each taste stimulus was lower than the taste- and temperature-selective neurons. We reasoned that this latter point could be the result of taste-selective only neurons being more narrowly tuned. Thus, to further investigate differences in the tuning profiles of taste-selective neurons, we computed the response sharpness index (SI), a standard technique used to evaluate the breadth of tuning of single neurons (Yoshida and Katz, 2011; Wilson and Lemon, 2013)(Fig. 8B), for each taste-selective neuron. High SI values are evidence of narrowly tuned neurons (i.e., GC neurons that encode 1 taste stimulus), whereas low SI values indicate broadly tuned neurons (i.e., GC neurons that encode multiple taste stimuli).

Figure 8B (*left*) shows the distribution of SI values for the two taste-selective groups. In agreement with our hypothesis, we observed that the taste- and temperature-selective neurons are more broadly tuned (Two Sample t-test: t = -2.0104, df = 111, p-value = 0.02341). These results were confirmed when we compared the fraction of taste-selective neurons having the SI included within a low (0*<*SI*>*0.35; 2-sample test for equality of proportions: X-squared = 4.765, df = 1, p-value = 0.01452) and a high (0.65*<*SI*>*1.0; X-squared = 2.9103, df = 1, p-value = 0.04401) range of values (Fig. 8B-*right*). These findings indicate that, with respect to their taste tuning profile, the taste- and temperature-selective neurons appear to be more broadly tuned, capable of being activated by more gustatory stimuli.

The above analysis examines only responsiveness, providing no direct information regarding whether the two groups of taste-selective neurons differ in how well they can encode taste information. We therefore performed a comparison of the distribution of the taste decoding accuracy for each of the taste-selective neurons (Fig. 8C-*left*). The decoding accuracy (computed with SVM, see Methods for more details) is a surrogate for the ability of each individual neuron to encode gustatory information in a 1.5-s post-stimulus temporal window. Visual examination of the data suggests that the spiking activity evoked by the gustatory stimuli in the taste- and temperature-selective neurons contains a greater amount of taste information. A non-paired *t* test confirmed the latter observation (Two Sample t-test: t = 1.7664, df = 111, p-value = 0.04004). This result is confirmed when comparing the fraction of taste-selective neurons with decoding accuracy *>* 0.55 (more than 2 ×chance-level)(Fig. 8C-*right* ; 2-sample test for equality of proportions: X-squared = 3.011, df = 1, p-value = 0.04135). Taken together, these observations indicated that GC neurons that were modulated by both chemosensory and thermal intraoral inputs were more effective in coding taste information than the GC neurons that responded only to gustatory inputs. We next sought to evaluate if the two populations of taste-selective neurons also differed in their capability of encoding information about taste palatability. Previous studies have indicated that GC neurons are capable of representing the palatability of intraoral taste stimuli (Katz et al., 2001; Bouaichi and Vincis, 2020) and that cooler temperatures of fluid solutions are preferred by water deprived rodents (Torregrossa et al., 2012). Based on these studies we hypothesized that the taste- and temperature-selective neurons would contain more palatability-related activity. We further reasoned that, if the latter hypothesis was confirmed by the data, it would shed some light in the direction of GC thermal responses representing mostly the reward - and less the sensory properties - of the intraoral thermal stimuli. However, contrary to our prediction, a two sample *t* test comparing the two distributions of PI revealed no differences between the taste- and temperature-selective and taste-selective only neurons (Fig. 8C; Two Sample t-test: t = -0.85211, df = 111, p-value = 0.198). Taken together, these observations indicate that taste- and temperature-selective neurons may represent - with respect to their taste responsive profile - a distinct set of broadly tuned neurons that can encode taste quality with high accuracy and may also participate in integrating taste quality with stimulus temperature.

## Discussion

This study evaluated the neural representations of intraoral thermal stimuli in the GC of behaving mice. We observed that GC neurons, as single units and as ensembles, were capable of reliably responding to and discriminating a wide range of innocuous intraoral temperatures of deionized water in a mostly monotonic manner. Intraoral responses appear to be distributed across the GC with the presence of an antero-postero gradient and a coarse dorso-ventral organization. Analyses of the similarity between the responses evoked by deionized water and artificial saliva revealed that, for the most part, temperature-related activity is driven by the thermal feature of the stimulus and not by the integration of the “taste of water” (Zocchi et al., 2017) and temperature. Finally our data indicate that thermal stimuli can recruit GC neurons that also respond to taste. Comparison of the response profiles of the GC neurons representing these two forms of intraoral stimuli and that of those being modulated exclusively by tastants revealed different taste quality coding and tuning properties, with taste- and temperature-selective neurons appearing more broadly tuned and capable of encoding more information than their taste-selective only counterparts; neither taste-selective group, however, seemed to encode more information about palatability than the other. Altogether, our findings demonstrate that the GC of behaving mice is involved in processing intraoral information important to ingestive behaviors. This work represents the first effort to reveal details of the cortical code for the mammalian intraoral thermosensory system in behaving mice and paves the way for future investigations on the cortical circuits and operational principles underlying thermogustation.

### Oral thermosensory coding in GC

Our results indicate that the GC integrates thermal information in the absence of an overt taste stimulus (Figs. 1-5). More than half of the GC neurons that were modulated by the contact of a 4-μl droplet of deionized water within the oral cavity were temperature-selective (Fig. 3B). It is important to note that some studies argue that water may be considered as an independent taste quality (Accolla et al., 2007) (Zocchi et al., 2017) and others have reported water-evoked responses along the gustatory neuraxis (Rosen et al., 2010; Bouaichi and Vincis, 2020). With this in mind, we compared the responses of neurons in the GC to different temperatures of deionized water and artificial saliva and found the evoked neural responses to be very similar. Indeed, a linear regression using deionized water at 14 °C and 36 °C as a stimulus significantly predicted the evoked firing rate for the same temperature of artificial saliva (Fig. 6D). We can, therefore, more confidently argue that the temperature-selective neurons recorded in the GC appear to encode mostly oral thermosensory information. The question that then arises is, what are the functions of the thermal intraoral signals that GC neurons may be representing? The first is stimulus identification, which involves sensory-discriminative processes aimed at qualitatively distinguishing between fluid temperature. When considering the temperatures used in our study in absolute terms, we observed GC responses to all three temperatures with often the same neurons capable of representing more than one stimulus (Fig. 3C). When considering the degree of thermal change (i.e., Δ*T*) achieved during stimulation from resting oral temperature [32 °C, (Leijon et al., 2019)], more GC neurons appear to be tuned to cooling (*<*32 °C) than to warming (*>*32 °C) stimuli. One challenge in the interpretation of this latter point is that we have not tested if our results generalize across a wider range of temperatures. For example, recent *in vivo* studies have shown that a subset of neurons in the trigeminal ganglion and the parabrachial nucleus of the pons (PBN) also respond to noxious cold (*<*14 °C) and hot (*>*40 °C) intraoral stimuli (Yarmolinsky et al., 2016; Lemon et al., 2016; Leijon et al., 2019; Li and Lemon, 2019; Lemon, 2021). The experimental design of the current work did not allow this level of analysis. We purposefully chose to omit noxious thermal stimuli to avoid distress for the mice and allow them to be engaged in the task and actively lick for a substantial number of trials. Another feature the GC may be encoding is the hedonic property of intraoral temperature. The thermal responses observed in our study could mostly reflect the hedonic - rather than the sensory quality - feature of the stimulus. Along this line, it is well established that GC neurons are capable of representing palatability of intraoral taste stimuli (Katz et al., 2001) and that cooler temperatures of fluid solutions are preferred by water deprived rodents (Torregrossa et al., 2012). As a result, one can expect a degree of convergence of taste and thermal stimuli onto palatability-related GC neurons. However, such an interpretation does not apply to our data. Indeed, GC neurons encoding both taste and temperature do not show a higher palatability index as compared to the ones encoding only gustatory stimuli (Fig. 8D).

Delving further into these taste-selective neurons, we show that intraoral thermal stimuli can modulate GC neurons that encode taste information (Figs. 7-8). Indeed, our results indicate that intraoral chemosensory and thermal inputs do not necessarily recruit distinct sets of GC neurons, but - for the most part - rather converge on the same cells. While this observation is in contrast with earlier reports (Kosar et al., 1986), it dovetails nicely with more recent studies highlighting the multimodal nature of many individual neurons in the GC (Vincis and Fontanini, 2016a; Samuelsen and Fontanini, 2017; Samuelsen and Vincis, 2021). Analysis of the taste-quality response profiles between GC neurons modulated by both intraoral stimuli (taste- and temperature-selective) and the ones encoding only taste information (taste-selective only) revealed some differences. While in both groups the majority of neurons were modulated by more than one taste quality (Fig. 8A), they differed in their breadth of tuning. Specifically, the GC neurons that were exclusively modulated by tastants (but not temperature; taste-selective only neurons) encoded fewer taste qualities (Fig. 8B), thus appearing to be more narrowly tuned. This observation is in agreement with a previous report showing that GC neurons modulated by retro-nasally delivered odor and taste are more broadly tuned to gustatory stimuli than the unimodal (i.e., modulated by taste only) neurons (Samuelsen and Fontanini, 2017). In addition, further analysis revealed that taste- and temperature-selective neurons encode taste quality - but not palatability-related - information with higher accuracy (Fig. 8). While we cannot yet provide definitive proof of the functional dissociation of the two groups of taste-selective neurons, we can speculate about their role in taste processing. GC neurons that encode both intraoral stimuli appear to be more broadly tuned within (taste quality) and across modality (taste and thermal inputs), and they may play a role in 1) the temperature-taste integration by linking thermal inputs to taste qualities - as well as in the “flavor network” (Small, 2012; Samuelsen and Fontanini, 2017; Samuelsen and Vincis, 2021) and 2) representing the neural substrate upon which associative flavor-related learning can operate (Vincis and Fontanini, 2016a). On the contrary, taste-only neurons may primarily encode one or two taste qualities in a way that is independent of their temperature or association with any other intraoral flavor-related sensory cue (Samuelsen and Fontanini, 2017).

### Spatial organization of orally-sourced thermal responses

Pioneering works in anesthetized rats have indicated that changes in intraoral temperature modulate the activity of a limited number of GC neurons (Kosar et al., 1986; Yamamoto et al., 1981). Interestingly, Kosar et al. provided some experimental evidence in support of a clear topographical organization of responses to intraoral stimuli. Their findings showed that, in the GC of anesthetized rats, thermal responses were not organized in a rostro-caudal fashion but rather appeared to be exclusively clustered dorsally (corresponding to the granular portion of the GC); taste responses, however, were clustered in the ventral agranular region (Kosar et al., 1986). To evaluate if intraoral thermal responses are topographically organized in the mouse GC, we performed a subset of electrophysiological recordings using linear and multi-shank silicon probes. With this approach, we relied on the high density of electrode contacts that allowed us to detect each spike at multiple sites, providing an opportunity to “triangulate” the location of each spike and to infer the relative 3D position of each single neuron. Our data revealed that temperature-selective activity along the dorso-ventral axis of the GC is mostly distributed with the exception of the most ventral portion of the GC, where there were more neurons responding to water in a temperature-independent manner (Fig. 5B). In addition, single neuron decoding revealed that the amount of temperature-selective information did not depend on the dorso-ventral position of the neuron (Fig. 5C). These findings appear to be partially in conflict with the one from Kosar et al. for at least two reasons. First, as already mentioned above, the distribution of temperature-selective neurons across the dorso-ventral axis in our study appears, for the most part, uniform. Second, our data indicate there is a high degree of convergence of taste and thermal responses in the same GC neuron (Fig. 8). This latter result - while in conflict with Kosar et al. where taste responses appeared to be exclusively clustered in the most ventral part of the rat’s GC - is in agreement with other studies in mice showing that spatial location plays no role in the distribution of taste responses (Levitan et al., 2019; Chen et al., 2021)[but see (Chen et al., 2011)]. It is important to highlight that the apparent lack of dorso-ventral topographical organization of thermosensory responses shown in our data does not necessarily exclude the possibility of the existence of subtle differences among GC subregions (granular, dysgranular, and agranular) or between the GC of different rodent models. Future experiments should endeavor to further probe these possibilities.

A recent study has elegantly shown that the posterior insular cortex - a cortical region rostral to the most posterior portion of the GC analyzed in this study - contains the primary cortical representation of skin temperature (Vestergaard et al., 2023). Interestingly, within the posterior insular cortex, body temperature appears to be topographically organized with thermal information from the paws being represented more posterior than the one originating from the face (Vestergaard et al., 2023). In our study, we investigated neural responses to intraoral thermal stimuli, focusing our attention on the GC that, as already mentioned above, is located rostral to the posterior insula region. While in our study, temperature-selective neurons encoded intraoral fluid with high accuracy irrespective of their antero-postero axis (Fig. 5E), there was a responsiveness gradient with more neurons being modulated by at least one temperature in the anterior part of the GC (Fig. 5D). It is thus tempting to speculate about the existence of a topographical organization of temperature in the broad insular cortex with a potential postero-antero gradient reflecting an extraoral to intraoral transition of thermal information.

### Origin of thermosensory responses in the GC

It is likely that one of the neural circuits allowing thermal information to reach the GC is in part the ascending gustatory pathway. Changes in intraoral temperature can stimulate somatosensory neurons of the trigeminal system, which are responsible for thermosensation of all oral surfaces (Lemon, 2021). *In vivo* experiments in mice have revealed that neurons in the trigeminal ganglia are highly sensitive to innocuous and noxious cooling (Yarmolinsky et al., 2016; Leijon et al., 2019; Lemon et al., 2016). Interestingly, while a subset of trigeminal ganglia neurons are also sensitive to noxious heat, comparably fewer cells respond to innocuous warming (Yarmolinsky et al., 2016; Leijon et al., 2019). In rodents, neurons in the trigeminal ganglia activate second-order trigeminal neurons that project to the PBN - a brain region that is also part of the ascending gustatory pathway (PBN receives taste inputs from the nucleus of the solitary tract). A recent study has discovered that some trigeminal inputs supplying craniofacial somatosensation reach the same PBN neurons receiving gustatory inputs (Li and Lemon, 2019). It is important to note that these trigeminal inputs reached PBN neurons that display sensitivity to aversive oral temperatures and tastes, highlighting the putative role of the PBN in processing and relaying multimodal intraoral sensory information including gustatory, nociceptive, and thermal stimuli (Li et al., 2022). Thermal stimulation inside the mouth can also recruit taste-sensitive nerves. On the one hand, temperature can influence the peripheral taste-sensitive neurons in presence of gustatory stimuli via the warmth-activated transient receptor potential (TRP) melastatin 5 (TRPM5) cation channel, which is involved with GPCR-mediated transduction cascades for sweet, umami, and bitter taste stimuli (Talavera et al., 2005; Zhang et al., 2003), and the thermal-sensitive epithelial sodium channels (ENaCs), which are involved with sodium taste transduction (Askwith et al., 2001). On the other hand, temperature can influence the peripheral taste-sensitive neurons even in absence of taste - a condition that is more relevant to the data presented in our study. Experimental evidence has indicated that a subset of chorda tympani neurons that do not respond to gustatory stimuli can be activated by cold fluid applied to the tip of the tongue (Yokota and Bradley, 2016, 2017). In addition, electrophysiological recordings of chorda tympani nerves in taste-deficient mice have revealed oral thermal responses (Finger et al., 2005). Future studies combining cell-type, molecular, and circuit manipulation approaches will be key to unraveling more details of the intraoral thermal pathway.

### A “taste” of things to come

In the past three decades, multiple studies have shed important light on the taste responses of GC neurons in awake rodents (Samuelsen and Vincis, 2021). Extracellular recordings and two-photon experiments have extensively and convincingly described both the temporal and spatial profile of taste-evoked activity. For example, electrophysiological data have highlighted the importance of hundreds of milliseconds- and seconds-long temporal dynamics of taste-evoked spiking activity (Katz et al., 2001; Gutierrez et al., 2010; Neese et al., 2022), while imaging experiments have revealed the lack of chemotopic organization of these responses (Chen et al., 2021)[but see also (Chen et al., 2011) and (Fletcher et al., 2017) for *in vivo* data]. However, these results originated from studies in which taste stimuli were experienced at a single temperature. While such an approach has shaped our understanding of cortical taste processing, it provides only a partial picture of the functional features of the GC. It is a common experience that temperature is a cue relevant to food preference, and multiple studies have shown temperature’s influence on taste perception (Moskowitz, 1973; Bartoshuk et al., 1982; Green and Frankmann, 1987; Torregrossa et al., 2012). In addition, our results indicate that GC activity is strongly modulated by intraoral fluid temperature and that gustatory and thermal inputs can converge on a subset of broadly tuned taste-selective neurons. For all of these reasons, one may wonder whether GC taste tuning, as well as the temporal and spatial properties of taste responses, are altered by the temperature of the stimulus and to what extent. Future experiments will examine the GC functional organization of chemosensory gustatory responses with respect to temperature-taste integration.

## Acknowledgments

This work was supported by funding from the National Institute of Deafness and Other Communication Disorders of the National Institute of Health Grant R01DC019326 to R.V. and Grant T32 DC000044 to C.G.B. The authors would also like to acknowledge Dr. Chris Lemon, Dr. Richard Bertram and the members of the Vincis laboratory for their feedback and insightful comments. We also thank Fred Fletcher and Te Tang for their excellent technical assistance. A portion of these data were presented in abstract form at the 2023 meeting of the Association for Chemoreception Sciences, Bonita Springs, FL.

## Author contribution

C.G.B. and R.V. designed research; C.G.B. and K.E.O. performed research; C.G.B., K.E.O., C.N. and R.V. analyzed data; R.V. wrote the first draft of the paper; K.E.O., C.G.B. and R.V. edited the paper; R.V. wrote the paper.

The authors declare no competing financial interest.

## Data availability

Zeonodo (data) and GitHub (code to reproduce figures and stats) accession numbers will be made available after acceptance.

